# A Deep Dive into Single-Cell RNA Sequencing Foundation Models

**DOI:** 10.1101/2023.10.19.563100

**Authors:** Rebecca Boiarsky, Nalini Singh, Alejandro Buendia, Gad Getz, David Sontag

## Abstract

Large-scale foundation models, which are pre-trained on massive, unlabeled datasets and subsequently fine-tuned on specific tasks, have recently achieved unparalleled success on a wide array of applications, including in healthcare and biology. In this paper, we explore two foundation models recently developed for single-cell RNA sequencing data, scBERT and scGPT. Focusing on the fine-tuning task of cell type annotation, we explore the relative performance of pre-trained models compared to a simple baseline, L1-regularized logistic regression, including in the few-shot setting. We perform ablation studies to understand whether pretraining improves model performance and to better understand the difficulty of the pre-training task in scBERT. Finally, using scBERT as an example, we demonstrate the potential sensitivity of fine-tuning to hyperparameter settings and parameter initializations. Taken together, our results highlight the importance of rigorously testing foundation models against well established baselines, establishing challenging fine-tuning tasks on which to benchmark foundation models, and performing deep introspection into the embeddings learned by the model in order to more effectively harness these models to transform single-cell data analysis. Code is available at https://github.com/clinicalml/sc-foundation-eval.

## 1 Introduction

Large-scale foundation models, which are pre-trained on massive, unlabeled datasets and subsequently fine-tuned on specific tasks, have recently achieved unparalleled success on a wide array of applications, including in healthcare and biology [1, 2, 3, 4, 5, 6]. The success of these models has showcased the power of leveraging generalizable features and contextual understanding to improve a model’s performance.

High-quality large-scale foundation models for single-cell RNA sequencing data could substantially improve the performance of single-cell RNA-sequencing analysis pipelines, and several such models have recently been developed, such as scBERT [7], scGPT [8], Geneformer [9], scFoundation [10], and scSimilarity [11]. The pre-train then fine-tune paradigm employed by these models holds the promise of allowing practitioners to leverage vast amounts of single-cell RNA sequencing data collected across a variety of technical and biological conditions, captured in pre-trained gene or cell embeddings. Fine-tuning the models on downstream tasks of interest then theoretically requires collection of much smaller, application-specific datasets than would be required if training a model from scratch. This sample efficiency is of particular value in the biological domain, where clinical data or data whose labels are determined experimentally can be challenging or expensive to collect, resulting in smaller available datasets for downstream tasks of interest.

In this paper, we explore two such models, scBERT [7] and scGPT [8]. Focusing on the fine-tuning task of cell type annotation, we provide further baselines and ablation studies beyond what was studied in the original papers, in order to examine the benefits of these foundation models for this task. We also explore the pretraining paradigm implemented by scBERT, and show that simple heuristics can achieve good performance at the pre-training task as it is currently formulated. Finally, using scBERT as an example, we demonstrate the potential sensitivity of fine-tuning to hyperparameter settings and parameter initializations. Taken together, our results highlight the importance of rigorously testing foundation models against well established baselines, establishing challenging fine-tuning tasks on which to benchmark foundation models, and performing deep introspection into the embeddings learned by the model in order to more effectively harness these models to transform single-cell data analysis.

## 2 Methods

### 2.1 Models Studied in this Paper

Figure 1 overviews the different models studied in this paper. The two foundation models studied in this work, scBERT and scGPT, rely on transformer-based architectures for processing embedded representations of the input gene expression data, but differ in how they represent input data, their model architectures, and training procedures. We also analyze a logistic regression baseline that directly trains on a specific task of interest without pre-training.

**Figure 1:**
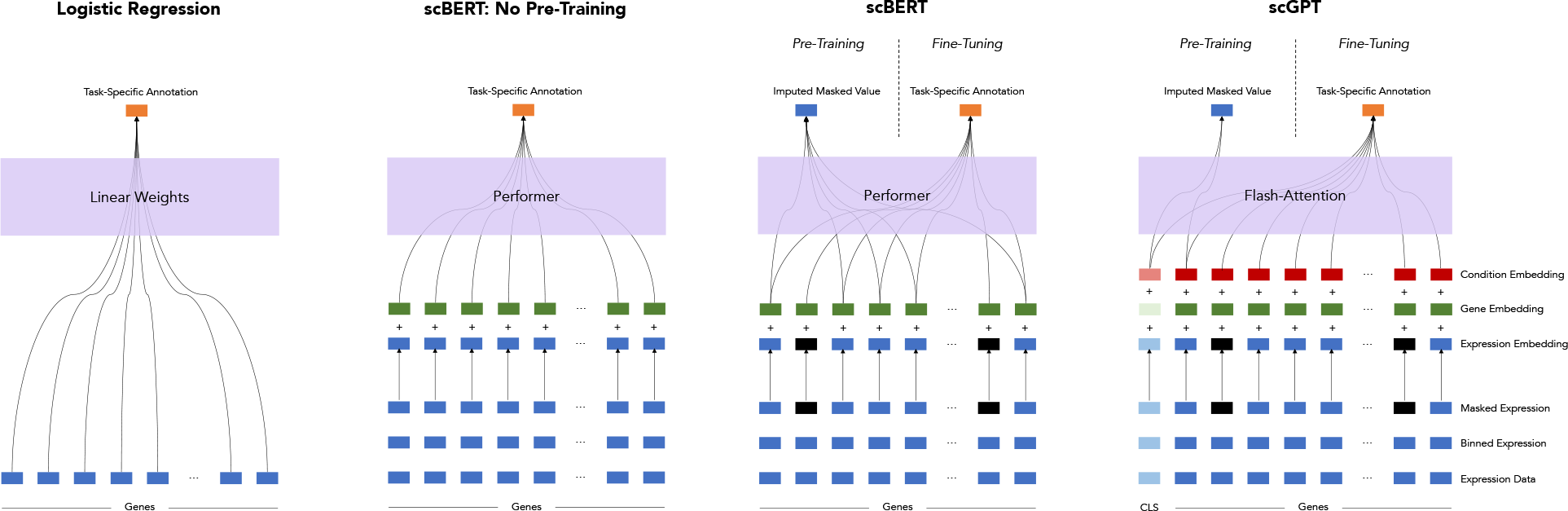
Schematic of different methods compared in this work. Two foundation models, scBERT and scGPT, construct embeddings of input data and process them via transformer-based architectures. The architectures are trained via a masking task and fine-tuned for a task of interest. We compare these models to logistic regression and a transformer architecture without pre-training to understand the potential benefits of the pre-train/fine-tune paradigm in several different experimental settings.

#### scBERT

scBERT [7] embeds each gene as the sum of two embeddings: one representing the gene’s normalized, binned log-scale expression level, and a second which specifies the gene identity via the previously developed gene2vec [12] embedding, allowing the model to differentiate genes in the otherwise order-less data. The resulting embeddings are processed via memory-efficient Performer [13] layers to produce the output.

Closely mimicking the pre-training strategy in BERT [5], scBERT is pre-trained via an imputation task on 1.12 million human cells where “masked” gene expression values are predicted as a function of all other gene embeddings in a cell (see Supplementary Methods). During fine-tuning, a 3-layer neural network is trained on smaller, labeled datasets to take the gene embeddings outputted by the transformer layers and predict cell type. In our analysis, we also evaluate a “no-pretraining” version of scBERT, in which pre-training is skipped, leaving the transformer layers frozen with randomly initialized weights at the time of fine-tuning.

#### scGPT

scGPT largely follows a similar architectural and pre-training paradigm to scBERT, with some different design choices. Instead of binning the input data on the log-scale, scGPT bins genes according to their expression such that genes are evenly distributed across each bin. In lieu of gene2vec, scGPT uses a random gene identity embedding and incorporates an additional “condition embedding” describing meta-information to differentiate each gene. Instead of the long-range Performer architecture, scGPT processes the embeddings via Flash-Attention [14] blocks. In addition to gene embeddings, scGPT further trains a CLS token which summarizes each cell. scGPT implements a *generative* masked pre-training, using a causal masking strategy inspired by OpenAI’s GPT series [15].

This model is pre-trained on 33 million human cells and evaluated via a wider suite of fine-tuning tasks, including cell type annotation, genetic perturbation prediction, batch correction, and multi-omic integration. We focus on cell type annotation in this paper.

#### Logistic Regression

Unlike the deep learning foundation models described previously, the logistic regression baseline [16] does not adhere to a pre-train/fine-tune paradigm. This method has the fewest number of trainable parameters and simply estimates the linear coefficients of the log-normalized gene expression data that best predict the labels of interest, in this case cell types. We use L1-regularization [17] to encourage sparsity.

### 2.2 Experiment Roadmap

In this work, we first explore the relative performance of pre-trained models like scGPT or scBERT compared to a simple baseline, L1-regularized logistic regression, for the fine-tuning task of cell type annotation. Since pre-trained representations can be especially useful for few-shot prediction, in which limited training data is available, we additionally evaluated each model’s ability to annotate cell types in the few-shot setting. We find that a simple logistic regression baseline is competitive with the pre-trained models for annotating cell types even in the few-shot setting.

Next, we sought to understand how much pre-training contributed to the performance of the transformer-based models. To this end, we skipped the pre-training procedure and directly fine-tuned the model for cell type annotation. We show that for scBERT, removing pre-training does not meaningfully affect the model’s performance on cell type annotation, while for scGPT, it does. We further demonstrate that scBERT can achieve good accuracy on masked pre-training and cell type annotation without learning meaningful gene representations, heeding that good accuracy does not necessarily imply rich representation learning.

Finally, we explore scBERT’s robustness to hyperparameter settings and random seeds, and we find that these settings significantly affect learning dynamics and model performance.

## 3 Results

### 3.1 Logistic regression outperforms foundation models for the fine-tuning task of cell type annotation in a dataset-dependent manner

To better understand scBERT’s performance on its main task, cell type annotation, we ran L1-regularized logistic regression [17] as a simple, interpretable baseline. Specifically, we predicted cell types from log-transformed, library size-normalized gene expression values in the Zheng68K peripheral blood mononuclear cells (PBMC) dataset [18], which contains 68,450 hand-labeled cells (80% used for training and 20% for validation; Supplementary Methods). This dataset was used for fine-tuning in the scBERT paper. Logistic regression outperformed scBERT in terms of both accuracy and class-averaged (macro) F1 score, a metric which reflects both precision and recall and can better assess performance in cases of class imbalance (Table 1). Even for difficult-to-distinguish CD8+ cytotoxic T cells and CD8+/CD45RA+ T cells [7], logistic regression outperformed scBERT in terms of accuracy and F1 score (Table 1).

**Table 1:**
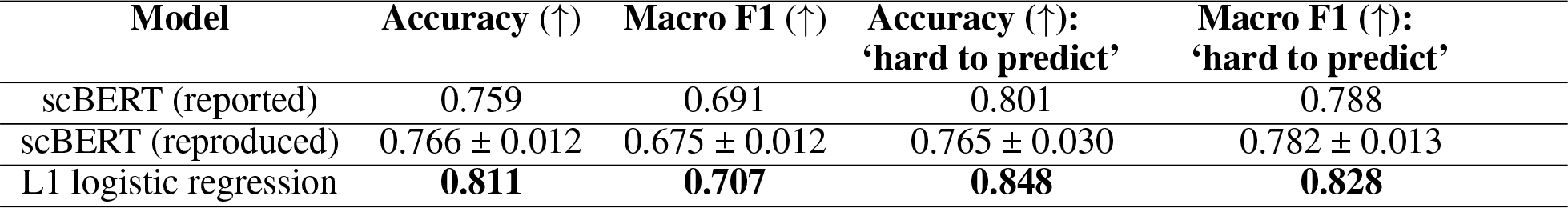
Logistic regression baseline for the cell type annotation task in scBERT. Validation set accuracy and macro F1 score for the cell type annotation fine-tuning task on the PBMC dataset, for three different models: scBERT as reported in the original publication [7], our reproduction of scBERT trained with the same data and model architecture, and our L1-regularized logistic regression baseline. scBERT was run with 10 different random parameter initializations and shufflings of the data during training; mean 95% confidence intervals reported. Logistic regression outperformed scBERT across all metrics for this task.

| Model | Accuracy (↑) | Macro F1 (↑) | Accuracy (↑):<br>'hard to predict' | Macro F1 (↑):<br>'hard to predict' |
| --- | --- | --- | --- | --- |
| scBERT (reported) | 0.759 | 0.691 | 0.801 | 0.788 |
| scBERT (reproduced) | $0.766 \pm 0.012$ | $0.675 \pm 0.012$ | $0.765 \pm 0.030$ | $0.782 \pm 0.013$ |
| L1 logistic regression | **0.811** | **0.707** | **0.848** | **0.828** |

Similarly, we ran logistic regression as a baseline for the cell type annotation task explored in the scGPT paper. The authors of scGPT evaluated the model’s cell type annotation capabilities on three different datasets: multiple sclerosis [19], pancreas [20], and myeloid data [21] (additional dataset details provided in Supplementary Methods). We found that for the multiple sclerosis data, scGPT outperformed logistic regression; for the myeloid data, logistic regression outperformed scGPT; and for the pancreas data, the two methods performed similarly (Figure 3; Supplementary Table 4).

These results show that classical baseline methods that are simpler to implement and less expensive to train may outperform or perform comparably to foundation models for certain predictive tasks on single-cell data.

### 3.2 Logistic regression can outperform foundation models even in the “few-shot” setting

A strong foundation model trains data representations that can easily adapt to downstream prediction tasks, even in “few-shot” settings with minimal training data [2]. Thus, while logistic regression outperformed pretrained models for cell type annotation in some datasets (PBMC, myeloid), we hypothesized that pre-trained models may show superior performance in the few-shot setting, as they can leverage the representations they learned from vast amounts of unlabeled single-cell data, while logistic regression only has access to the small set of training examples.

We found, however, that across models, datasets, and fractions of training data used, the trends from Section 3.1 generally hold even in the few-shot setting, with a few exceptions. In particular, as in Section 3.1, logistic regression outperforms or matches the foundation models on the PBMC and myeloid data even with limited training data (Figures 2, 3). Similarly, just as in Section 3.1, scGPT outperformed logistic regression on the multiple sclerosis data across all training set sizes, with the relative benefit of the pre-trained model increasing with decreasing training set size (Figure 3). For the pancreas dataset, logistic regression and scGPT were closely matched with larger training set sizes, but logistic regression outperformed scGPT once the training set was limited to just 10% or 25% of the original training set size (Figure 3). Detailed performance metrics and two-tailed t-test p-values describing these comparisons are shown in Figures 2 and 3.

**Figure 2:**
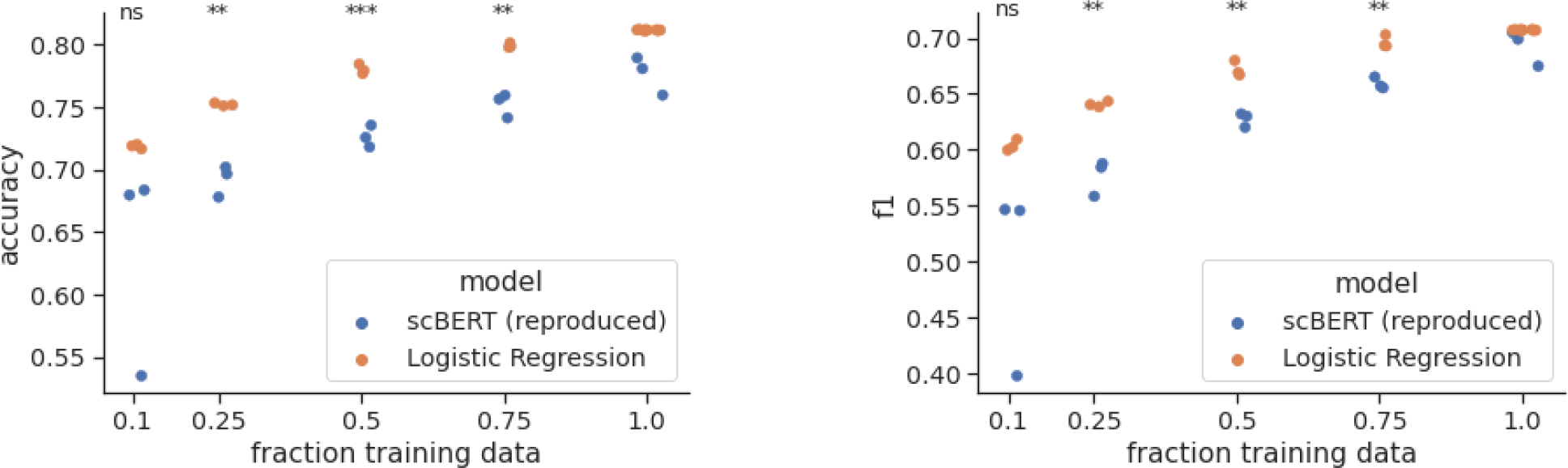
scBERT and logistic regression validation set performance (accuracy, left; F1 score, right) for the fine-tuning task of cell type annotation in the PBMC dataset, using different amounts of training data (jitter added to x-axis to better display overlapping data points). The same pre-trained model was used in all scBERT runs. Each experiment was run independently 3 times, each time randomly subsetting the training set and, for scBERT, randomly initializing model parameters and shuffling the order of the training data. Logistic regression consistently outperformed scBERT even with small amounts of training data. A two-sided t-test was used to compare performance of the models for each amount of training data: *** *p <*= 0.001; ** *p <*= 0.01; “ns” not significant (*p >* 0.05).

**Figure 3:**
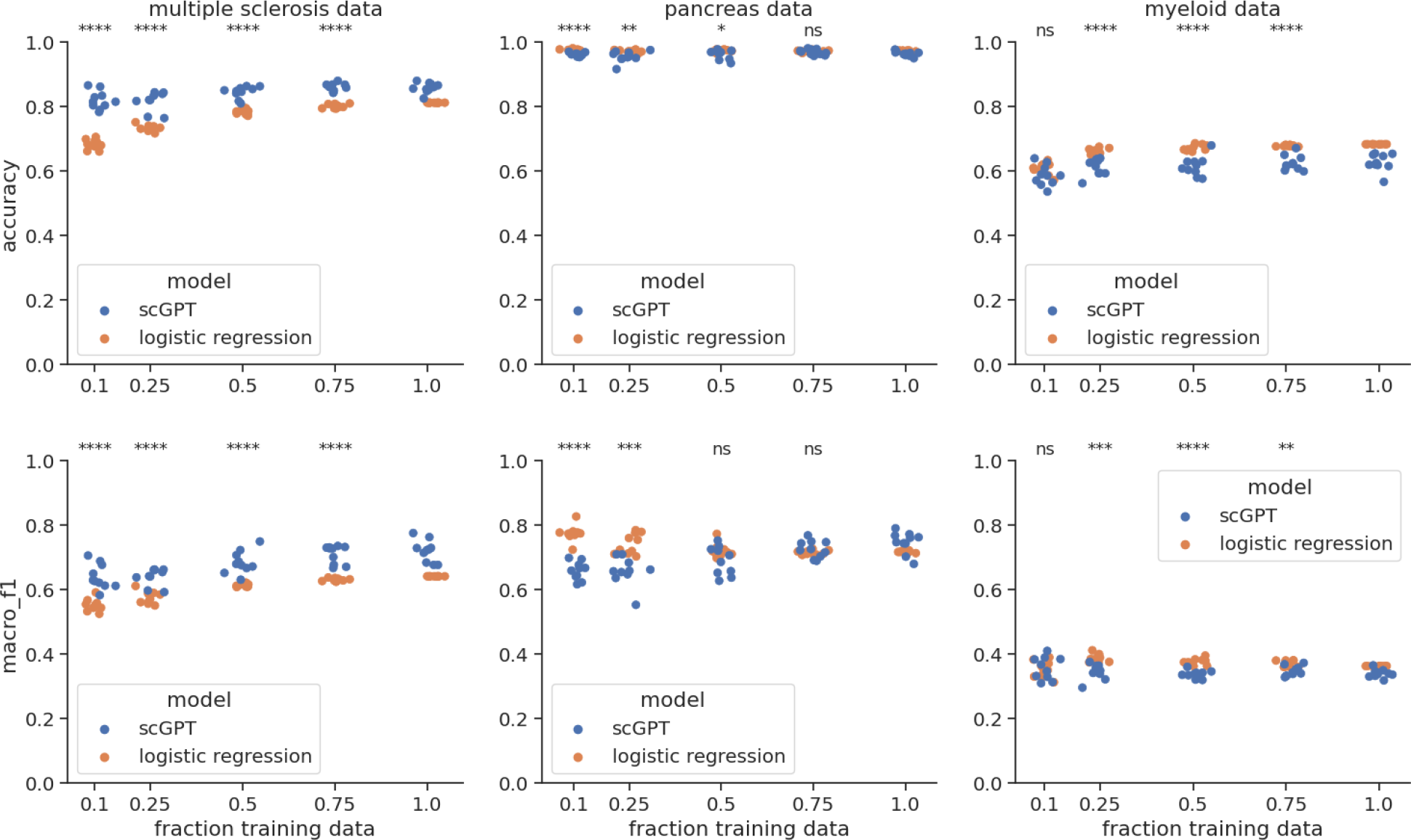
scGPT and logistic regression test set performance (accuracy, top; F1 score, bottom) for the fine-tuning task of cell type annotation in the three datasets used in the scGPT paper, using different amounts of training data (jitter added to x-axis to better display overlapping data points). The same pre-trained model was used in all scGPT runs. Each experiment was run 10 times with different random seeds, each time randomly subsetting the training set and, for scGPT, randomly initializing model parameters. For each dataset, a two-sided t-test was used to compare performance of the models for each amount of training data: **** *p <*= 0.0001; *** *p <*= 0.001; ** *p <*= 0.01; * *p <*= 0.05; “ns” not significant (*p >* 0.05).

One possible explanation for the different performance trends across these datasets lies in the different nature of the datasets. Among all the datasets, the pretrained model specifically excelled with the multiple sclerosis dataset, which was crafted by the scGPT authors to represent an “out of distribution” scenario: the fine-tuning training set consisted of neuronal cells from 9 healthy donors, and the test set consisted of neuronal cells from 12 multiple sclerosis patients. The pre-trained model may be better equipped to handle prediction under dataset shift, having been pre-trained on a vast number of cells from across many different clinical contexts, whereas logistic regression has only been trained on a single clinical context that is no longer relevant in the test set. In other words, pre-training may serve as a form of regularization that protects the model from overfitting to the training distribution and helps it to generalize.

None of the other datasets studied in this work suffer from as drastic a distribution shift. Thus, in these scenarios, the cost of training a high parameter model with a small amount of training data may outweigh the benefit of pre-training, and thus logistic regression outperforms or performs comparably to scGPT in the low-data regime. Analyzing the relative performances of the pre-trained and logistic regression models across these different data settings suggests that pre-trained models have unique strengths that make them the better choice for some, but not all, single cell predictive tasks.

### 3.3 Skipping pre-training does not affect fine-tuning performance for scBERT, but does for scGPT

The paradigm of “pre-train then fine-tune” assumes that the representations learned during unsupervised pre-training help the model perform well on downstream tasks. To test this, we ran an ablation study in which we skipped the pre-training step, effectively using a random embedding of a cell’s expression data as input for predicting cell type, to evaluate whether cell type annotation performance decreased without pre-training (see Supplementary Methods).

For scBERT, we found that removing pre-training did not significantly change performance on annotating PBMC cell types (Table 2; two-tailed t-test *p* = 0.34 and *p* = 0.59 for accuracy and F1 score, respectively). This finding may indicate that the scBERT pre-training task does not meaningfully improve the representations learned by the model, or may reflect the simplicity of the cell type annotation task for PBMC data.

**Table 2:** Ablation studies for the cell type annotation task in scBERT. Accuracy and macro F1 score for the cell type annotation fine-tuning task on the PBMC dataset, for three different models: our reproduction of scBERT trained with the same data and model, and ablations that remove the pre-training step and gene2vec embedding, respectively. All results are reported on the validation set. Each model was run with 10 different random parameter initializations and shufflings of the data during training; mean performance *±*95% confidence intervals across these runs are shown. We found that without pre-training, none of the performance metrics significantly changed. Without gene2vec embeddings, accuracy and F1 averaged across all cell types decreased (two-tailed t-test *p* = 3.67 *×* 10^*−*3^ and 8.72 *×* 10^*−*4^, respectively) and F1 for the “hard to predict” cell types decreased (two-tailed t-test *p* = 4.10 *×* 10^*−*3^).

| Model | Accuracy (↑) | Macro F1 (↑) | Accuracy (↑):<br>‘hard to predict’ | Macro F1 (↑):<br>‘hard to predict’ |
| --- | --- | --- | --- | --- |
| scBERT (reproduced) | $0.766 \pm 0.012$ | $0.675 \pm 0.012$ | $0.765 \pm 0.030$ | $0.782 \pm 0.013$ |
| No pre-training | $0.758 \pm 0.014$ | $0.672 \pm 0.006$ | $0.754 \pm 0.035$ | $0.772 \pm 0.016$ |
| No gene2vec | $0.701 \pm 0.040^{**}$ | $0.595 \pm 0.042^{**}$ | $0.714 \pm 0.046$ | $0.709 \pm 0.046^{**}$ |

By contrast, the scGPT authors report results from a similar experiment, “scGPT (from-scratch),” in their Supplementary Table S1, finding that their pre-trained model outperformed similar models fine-tuned “from-scratch,” without pre-training. We independently ran our “no-pretraining” ablation on scGPT, and also concluded that pre-training improved performance on cell type annotation for scGPT across all three datasets (Supplementary Table 5 and Supplementary Figure 6).

Further analysis is needed to understand whether the difference in “no pre-training” performance between scBERT and scGPT can be ascribed to the different models, or the different fine-tuning datasets evaluated.

### 3.4 For scBERT’s embedding scheme and pre-training objective, good pre-training and fine-tuning accuracy can be achieved without learning rich representations

Despite good performance on the pre-training objective (78% accuracy), Section 3.3 shows that pre-training scBERT does not affect the final fine-tuning performance. In this section, we ablate another component of the scBERT architecture and show that good pre-training or fine-tuning accuracy is not sufficient evidence that a model is learning rich gene representations.

In scBERT, a given gene’s embedding is the sum of two separate embeddings for that gene: a “gene2vec” embedding and an “expression” embedding. The “gene2vec” embedding codifies the identity of each gene and is constant for each gene across every cell in the dataset. The “expression” embedding transforms a given gene’s expression into one of six randomly-initialized vectors based on its binned expression (the bins are [0-1), [1-3), [3-7), [7-15), [15-31), and 31+ library size-normalized UMIs). The expression embedding – not the gene2vec embedding – drives the variability in the learned representations of a given gene for different cells and is the focus of the masking task used to pre-train the model.

In this experiment, we remove the gene2vec embedding from the representation of each gene. This severely limits the representational capacity of the model, as there are now only six embeddings that are used to represent every gene in the dataset. During pre-training, the model has no knowledge of a gene’s identity and every masked gene has identical “context” (note that since transformers are permutation invariant [22], the model cannot memorize the input positions of genes in order to identify them). Every gene that falls in the same expression bin is represented identically – for example, across the Panglao pre-training dataset [23], 95% of genes are identically represented as falling in the [0,1) bin, 2% of genes in the [1-3) bin, 1.5% of genes in the [3-7) bin, and <1% of genes in the remaining bins.

Given that the model does not know which gene is which, one might expect that scBERT would not be able to predict a gene’s masked expression with high accuracy. We observed, however, that even in this challenging setting, scBERT achieved 72% accuracy for predicting masked gene expression on the Panglao validation set (compared to 78% when genes are represented using both gene2vec + expression embeddings). Upon introspection into the “no gene2vec” experiment, we found that for 99.8% of the 67,880 cells in the Panglao pre-training validation set, scBERT predicted the most common expression bin per cell for all masked genes in that cell (Table 3). These results demonstrate that with scBERT’s binned expression embedding scheme, a model can rely on simple heuristics such as predicting the most common expression bin to perform well at masked pre-training without learning contextual gene representations.

**Table 3:** Validation set accuracy on masked pre-training when gene2vec embeddings are included (“full scBERT”) or are not included (“no gene2vec”) as part of each gene’s input embedding. In the no gene2vec setup, the same expression bin is necessarily predicted for all masked genes in a given cell, as they all have identical context. This is not the case in the full scBERT embedding scheme; hence “full scBERT” has N/A in the column quantifying the fraction of cells for which the most common expression bin is predicted, as only gene-level predictions can be summarized in this way.

| Model | Data | Pre-training<br>masking accu-<br>racy (↑) | Fraction of cells in which<br>most common expression bin<br>was predicted for all genes | Fraction of masked genes<br>for which the most common<br>bin (per cell) was predicted |
| --- | --- | --- | --- | --- |
| no gene2vec | PBMC | 68% | 100% | 100% |
| full scBERT | PBMC | 79% | N/A | 79% |
| no gene2vec | Panglao | 72% | 99.8% | 99.9% |
| full scBERT | Panglao | 78% | N/A | 84% |

Regarding fine-tuning, scBERT trained without gene2vec embeddings exhibited decreased but still reasonable performance on the cell type annotation task (Table 2, “no gene2vec” accuracy 0.701 and F1 0.595). We note that we did not re-tune hyperparameters specifically for the no gene2vec experiment, but rather used the best hyperparameters we identified for fine-tuning the full scBERT model, so it is possible that the performance we report here is an underestimate of the potential performance had we tuned hyperparameters for this task specifically (see Section 3.5). While the logistic regression baseline and no-pretraining ablation already demonstrated that rich representations are not needed for accurate cell type annotation in the PBMC data, this result further reinforces that performance on cell type annotation is not a proxy for the representation learning capabilities of a foundation model.

### 3.5 Robustness to hyperparameter choices and parameter initialization

In working to reproduce scBERT’s reported accuracy for cell type annotation in the PBMC data, we found that batch size and learning rate - neither of which were reported in the scBERT paper - were critical hyperparameters to tune in order to achieve optimal fine-tuning results. Adopting a learning rate of 1 *×* 10^*−*4^, as was the default setting in the scBERT code base, we evaluated validation set performance across multiple batch size settings, and observed that smaller batch sizes, such as 8 or 32, generally achieved better performance (Figure 4). We chose to use a batch size of 32 in all our experiments reported in this paper, to strike a balance between efficiency (large batch sizes are more efficient) and optimization (smaller batch sizes achieve more optimal results).

**Figure 4:**
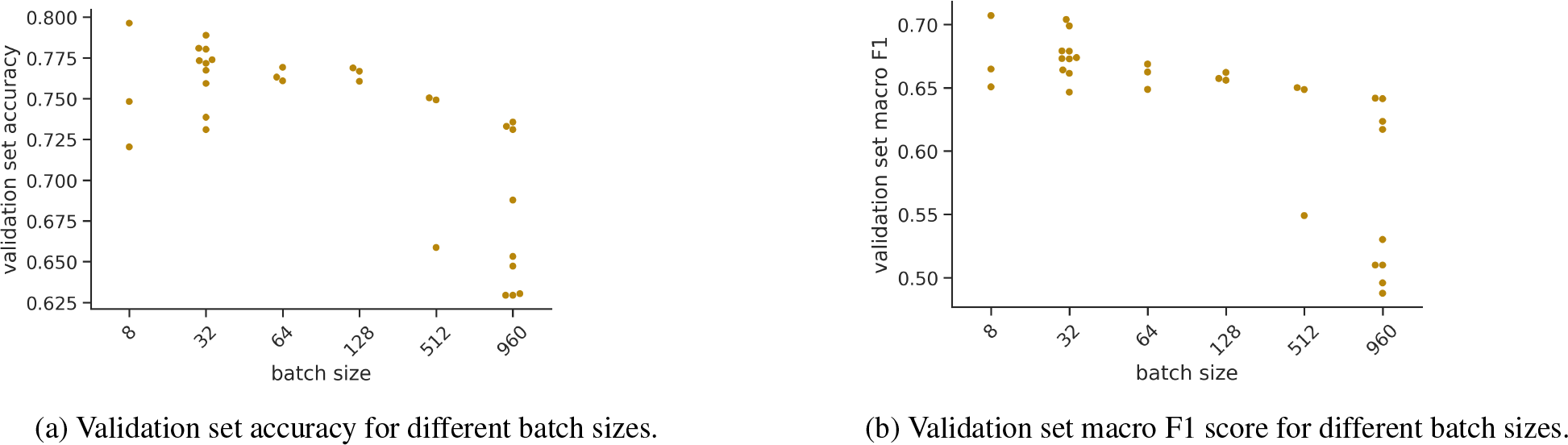
The mini-batch size used for fine-tuning scBERT for the cell type annotation task has a large effect on predictive performance, in terms of validation set accuracy (a) and macro F1 score (b). All models shown here were trained with a learning rate of 1 *×* 10^*−*4^. Each model was trained for 10 epochs, and metrics from the epoch with the best validation set accuracy are shown. Each hyperparameter setting was run multiple times with different random seeds, each represented by a single dot.

In addition to batch size, we explored the effect of random seed (i.e. parameter initialization) on predictive performance. The random seed determines the initial random parameterization of the fine-tuning components of the scBERT architecture (these include a convolutional filter to summarize each gene embedding, and a 3-layer neural network). Similar to results reported by Liu et al. [24], we found that performance indeed varied across random seeds, and that fluctuations in performance across random seeds were greater for models trained with larger batch sizes (Figure 4). Even for an optimal batch size of 32, we observed that two different learning patterns emerged across the random seeds: one group of random seeds led to convergence in fewer epochs and achieved overall higher accuracy after ten training epochs, while a second group of random seeds led to longer convergence times and overall lower accuracy after ten training epochs (Figure 5). Understanding this bimodal behavior of the learning dynamics requires further exploration. We note that if the models with low-performing initializations were trained for additional epochs, they may eventually reach the quality of the high-performing ones.

**Figure 5:**
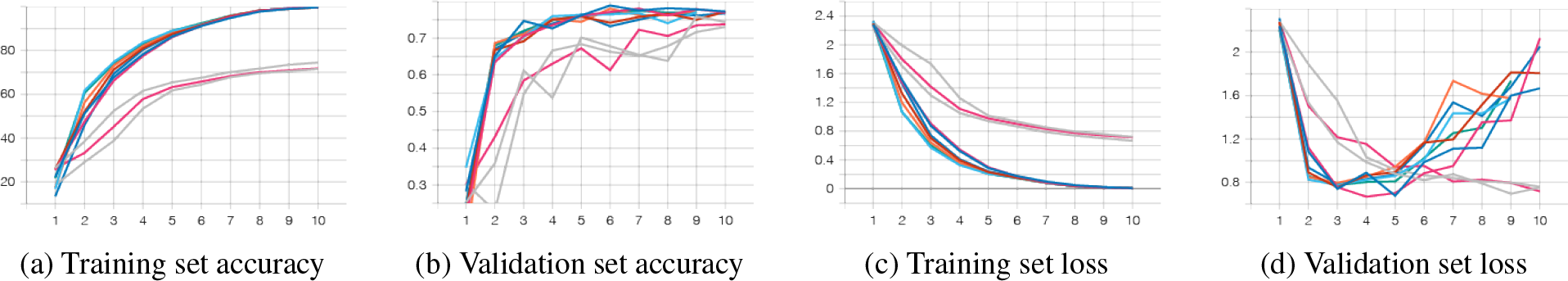
Accuracy and loss per epoch for fine-tuning scBERT for cell type annotation, for ten different models initialized with different random seeds, but otherwise identical. All models were trained with batch size 32 and learning rate 1 *×* 10^*−*4^ for 10 epochs. We observe that parameter initialization affects both learning dynamics and final performance metrics.

These findings emphasize that in order for a model to be reproducible, 1) a full set of hyperparameter settings must be transparently reported and 2) performance should be reported over multiple random seeds, as we did in Table 1 and Supplementary Table 4. While releasing code is incredibly helpful and important, it is not sufficient to only specify hyperparameters as default settings in a code base, as it is all too easy for users to inadvertently change hyperparameter settings when running code in a new compute environment (eg. training with the same batch size setting on 1 GPU vs. 4 GPUs in a distributed fashion may result in batch size effectively being scaled by 4).

## 4 Discussion

As interest in foundation models for single-cell RNA sequencing data grows within the genomics community [9, 8, 11, 10, 24], we aimed to provide a deeper understanding of their potential benefits and limitations by providing additional baselines and introspection into the models’ components.

We demonstrated that simple models like logistic regression can be competitive for the fine-tuning task of cell type annotation, even in the few-shot setting. We further showed that even without pre-training, transformer architectures can perform well at cell type annotation. These results underscore that cell type annotation is not a challenging enough task on which to demonstrate the potential value of single-cell foundation models. The value of foundation models for single-cell RNA sequencing data will instead be showcased through superior performance at more challenging fine-tuning tasks which simple models currently cannot solve, or else through the models’ abilities to capture and elucidate meaningful gene-gene interactions through their learned attention weights. When claiming the latter, it is important to keep in mind that strong downstream performance does not necessarily imply rich representation learning, as shown in Section 3.4. Finally, since hyperparameter settings and the initial parameterization of models weights can have large effects on the learned model, it is important to have transparent reporting of all design choices and hyperparameter settings, in addition to sharing code and trained models, in order for the community to reproduce and build off of each other’s works.

Taken together, we hope that these results shed light on the importance of deep introspection into model performance, as well as rigorous baselines and evaluation tasks, to demonstrate the value and justify the compute costs of large scale foundation models for RNA sequencing data. We hope that these findings will enable and encourage the community to improve on current transformer-style architectures for single-cell RNA sequencing data.

## 5 Supplementary Data

**Table 4:** A comparison of logistic regression and scGPT for cell type annotation. Accuracy and macro F1 scores for the cell type annotation fine-tuning task on the three datasets evaluated in the scGPT paper. Each deep model was run with 10 different random parameter initializations and shufflings of the data during training; mean performance *±* 95% confidence intervals across these runs are shown. The best metric for each dataset is bolded.

| Dataset name | Model | Test set accuracy | Test set macro F1 score |
| --- | --- | --- | --- |
| multiple sclerosis | logistic regression | 0.812 | 0.642 |
| multiple sclerosis | scGPT | **0.859 ± 0.010** | **0.720 ± 0.023** |
| pancreas | logistic regression | **0.973** | 0.718 |
| pancreas | scGPT | 0.965 ± 0.005 | **0.748 ± 0.023** |
| myeloid | logistic regression | **0.683** | **0.364** |
| myeloid | scGPT | 0.627 ± 0.018 | 0.340 ± 0.008 |

**Figure 6:**
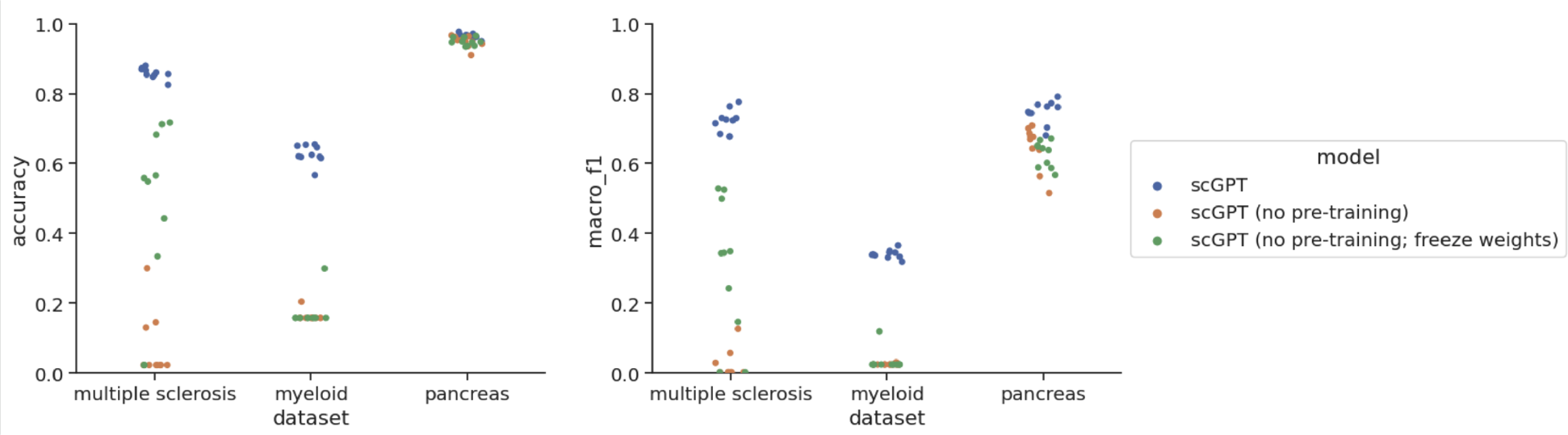
Accuracy (left) and macro F1 score (right) for cell type annotation on 3 datasets for the no pre-training ablation in scGPT (with and without freezing weights) vs. vanilla scGPT fine-tuning. Each model was run with 10 different random seed initializations. Mean and 95% confidence intervals for each model are shown in Supplementary Table 5. We note that while we reach the same conclusion as Cui et al. [8] that pre-training benefits downstream performance for cell type annotation, these results significantly differ from what was reported in their Supplementary Table S1; further exploration is needed to understand the source of this discrepancy.

**Table 5:** Cell type annotation results when skipping pre-training for scGPT. Accuracy and macro F1 scores for the cell type annotation fine-tuning task on the three datasets evaluated in the scGPT paper for three different models: scGPT (with pre-training), and two different versions of an ablation skipping pre-training, one in which all weights of the model were learnable during fine-tuning, and one in which all pre-decoder weights were frozen during fine-tuning. Each model was run with 10 different random parameter initializations and shufflings of the data during training; mean performance *±* 95% confidence intervals across these runs are shown. The best metric for each dataset is bolded. We note that while we reach the same conclusion as Cui et al. [8] that pre-training benefits downstream performance for cell type annotation, these results significantly differ from what was reported in their Supplementary Table S1; further exploration is needed to understand the source of this discrepancy.

| dataset_name | model | accuracy | macro_f1 |
| --- | --- | --- | --- |
| multiple sclerosis | scGPT | **0.859 ± 0.01** | **0.72 ± 0.023** |
| | scGPT (no pre-training) | 0.074 ± 0.063 | 0.023 ± 0.028 |
| | scGPT (no pre-training; freeze weights) | 0.461 ± 0.176 | 0.298 ± 0.134 |
| myeloid | scGPT | **0.627 ± 0.018** | **0.34 ± 0.008** |
| | scGPT (no pre-training) | 0.163 ± 0.01 | 0.025 ± 0.001 |
| | scGPT (no pre-training; freeze weights) | 0.172 ± 0.03 | 0.034 ± 0.02 |
| pancreas | scGPT | **0.965 ± 0.005** | **0.748 ± 0.023** |
| | scGPT (no pre-training) | 0.949 ± 0.011 | 0.648 ± 0.042 |
| | scGPT (no pre-training; freeze weights) | 0.949 ± 0.008 | 0.627 ± 0.025 |

## 6 Supplementary Methods

### 6.1 Additional dataset details

#### Multiple Sclerosis

The multiple sclerosis data was comprised of neuronal cell types from healthy donors and patients with multiple sclerosis. We downloaded the pre-processed data provided by the scGPT authors, thus using the same pre-processing and train/test splits as they did. The train/test split for this dataset simulates dataset shift, with cells from 9 healthy donors comprising the training set, and from 12 multiple sclerosis patients comprising the test set. There are 7,844 cell in the training set and 13,468 in the test set. The ground truth cell-type labels were taken from the original publication of the data [19].

#### Myeloid

This dataset contains tumor infiltrating myeloid cells from multiple different tumor types [21]. We downloaded the pre-processed data provided by the scGPT authors, thus using the same pre-processing and train/test splits as they did. In particular, the dataset consisted of cells from nine different cancer types, with the train set containing cells from cancer types UCEC, PAAD, THCA, LYM, cDC2, and KIDNEY, and the test set containing cells from MYE, OV-FTC, and ESCA. The final cell counts are 9,748 in the training set and 3,430 in the test set.

#### Pancreas

The pancreas dataset contained human pancreas cells from five different scRNA-seq studies, which were reprocessed by Chen at al. [20]. The training set consisted of 10,600 cells of 13 cell types (alpha, beta, ductal, 808 acinar, delta, PSC, PP, endothelial, macrophage, mast, epsilon, schwann, and t cell) from two datasets, and the test set consisted of 4,218 cells of 11 cell types (alpha, beta, ductal, PP, acinar, delta, PSC, endothelial, 810 epsilon, mast, MHC class II) from the other three datasets. We downloaded the pre-processed data provided by the scGPT authors, thus using the same pre-processing and train/test splits as they did.

### 6.2 Pre-training and fine-tuning scBERT

In the original scBERT paper, the scBERT model was pre-trained on the Panglao dataset provided by the scBERT authors, which is comprised of gene expression from multiple studies spanning 74 human tissues. The model was then fine-tuned on individual smaller datasets, for example the Zheng68k PBMC dataset, and metrics were reported for the performance of the model in annotating cell types in the fine-tune dataset. In our experiments, we ran pre-training on the Panglao dataset (provided by the original scBERT authors upon request, with 95% of the data used for training and 5% for validation). We masked the expression embedding for 15% of genes in a given cell (i.e. set the embedding to a randomly-initialized “mask” embedding). Genes with log-normalized UMIs between [0,1) were excluded from the masking task in scBERT in order to avoid an overwhelming class imbalance, since >95% of the expression values in the Panglao dataset were zero. We pre-trained the model using the Adam optimizer [25] in PyTorch, with a batch size of 8, gradient accumulation every 60 steps and a learning rate of 1 *×* 10^*−*4^ for 17 epochs.

After pre-training the model, we ran fine-tuning on the Zheng68K PBMC dataset (80% training, 20% validation), using the scripts provided by the authors, with minimal changes. For fine-tuning, we fixed the training and validation sets to be constant across different runs and models, so that our evaluations would isolate differences in performances due to changes in the model, rather than changes in the training or validation data. We trained all fine-tuning experiments using the Adam optimizer [25] in PyTorch with a learning rate of 1 *×* 10^*−*4^ and batch size of 32 (with no gradient accumulation) for 10 training epochs, and reported results from the epoch corresponding to the best validation set accuracy.

For pre-training and fine-tuning, we slightly modified the scBERT authors’ provided code to use pyTorch’s DistributedDataParallel module (https://pytorch.org/tutorials/intermediate/ddp_tutorial.html) to allow us to train over multiple GPUs in our compute environment. For fine-tuning, we further adjusted the code to weight the loss for different classes differently in order to counter class imbalance and improve the model’s ability to generalize outside the training data, and we modified the dataloader to load the same training samples in each epoch of training (shuffling their order), whereas the original scBERT code used a different sampling of the training data in each epoch.

### 6.3 Fine-tuning scGPT

We followed the fine-tuning procedure outlined in the original scGPT paper and implemented in the tutorial provided by the authors at https://github.com/bowang-lab/scGPT/blob/main/tutorials/Tutorial_Annotation.ipynb. For few-shot experiments, we randomly subsampled the training datasets in a stratified fashion to ensure that the same class balance was present in the full and subsampled training data. We used all default settings and hyperparameters provided in the authors’ code, including a learning rate of 1 *×* 10^*−*4^ and batch size of 32. For cell type annotation in the full-data and few-shot settings, we fine-tuned for 20 training epochs and selected the model from the epoch with the best validation set performance for reporting test set results. For fine-tuning the “no pre-training” experiment, we fine-tuned for 30 training epochs, in case further training was needed to compensate for the lack of pre-trained embeddings.

### 6.4 Logistic Regression

For our logistic regression baseline, we split the data into an 80:20 train:test split and performed 5-fold cross validation using the training data to choose the regularization coefficient. For experiments that used 100% of the training data, the following regularization coefficients (“c” in sklearn’s sklearn.linear_model.LogisticRegression function) were chosen:

- PBMC: c=0.1
- multiple sclerosis: c=0.1
- pancreas: c=100
- myeloid: c=0.1

For the few-shot experiments, a different c was chosen using cross validation for each subsample of the training data.

### 6.5 Few-shot experiments

To evaluate performance with limited training data, we trained each model on 75%, 50%, 25%, and 10% of its full training data. We used the same train/test split as in our other fine-tuning experiments, randomly sub-sampling a fraction of the original training data. For each new subsample of training data, we performed 5-fold cross validation using the smaller training data to choose the regularization coefficient. We evaluated performance on the full test set. We ran each experiment 3 times for the PBMC data and 10 times for the multiple sclerosis, pancreas, and myeloid datasets, each time using a different random sub-sample of the training data and a different random seed to initialize parameters, to capture the variation resulting from both the composition of the small training set and the model’s initial parameter settings.

### 6.6 “No pre-training” ablation

For the “no pre-training” ablation experiment on scBERT, we randomly initialized the full scBERT architecture, but instead of pre-training, we directly fine-tuned this model on the PBMC data for the cell type annotation task. Similar to our other runs of pre-training, we froze most of the (random, in this case) weights of the transformer, leaving only the weights in the last layer of the transformer and in the fine-tuning portions of the architecture to be updated during fine-tuning. We fine-tuned each model for 10 epochs, and reported results for the epoch with the highest validation set accuracy.

For the “no pre-training” ablation experiment on scGPT, we took a very similar approach, randomly initializing the full scGPT architecture and directly fine-tuning the model for cell type annotation on either the myeloid, multiple sclerosis, or pancreas dataset without learning or loading any pre-trained weights onto the model. We experimented with two different modes of this experiment, one in which all weights in the model were trained during fine-tuning, and one in which all pre-decoder weights were frozen, reducing the number of trainable parameters from 51,342,358 to 19,999,766. We fine-tuned each model for 30 epochs, and reported results for the epoch with the highest validation set accuracy.

## 7 Data Availability

All data used in this manuscript can be accessed using the links provided by the scBERT and scGPT authors. In particular, the Panglao dataset can be accessed at https://drive.google.com/file/d/1fNZbKx6LPeoS0hbVYJFI8jlDlNctZxlU/view?usp=sharing, and the Zheng68K PBMC dataset and pre-trained gene2vec embeddings can be accessed at https://drive.weixin.qq.com/s?k=AJEAIQdfAAozQt5B8k. The myeloid, multiple sclerosis, and pancreas datasets can be accessed at https://github.com/bowang-lab/scGPT/tree/main/data.

## 8 Code Availability

Our scripts and analysis code are available at https://github.com/clinicalml/sc-foundation-eval.

## 9 Acknowledgements

We thank Ava Amini for her feedback on our experiments and manuscript draft. R.B. and D.S were supported by an ASPIRE award from The Mark Foundation for Cancer Research. G.G. is partially supported by the Paul C. Zamecnik Chair in Oncology at the Massachusetts General Hospital Cancer Center. N.S. was funded by a PhD fellowship from Google.

